# Viability of MS2 and Phi6 Bacteriophages on Carpet and Dust

**DOI:** 10.1101/2021.05.17.444479

**Authors:** Nicholas Nastasi, Nicole Renninger, Ashleigh Bope, Samuel J. Cochran, Justin Greaves, Sarah R. Haines, Neeraja Balasubrahmaniam, Katelyn Stuart, Jenny Panescu, Kyle Bibby, Natalie M. Hull, Karen C. Dannemiller

**Affiliations:** Environmental Sciences Graduate Program, Ohio State University, Columbus, OH 43210; Department of Civil, Environmental & Geodetic Engineering, College of Engineering, Ohio State University, Columbus, OH 43210; Division of Environmental Health Sciences, College of Public Health, Ohio State University, Columbus, OH 43210; Department of Civil & Environmental Engineering & Earth Sciences, College of Engineering, University of Notre Dame, Notre Dame, IN 46556; Sustainability Institute, Ohio State University, Columbus, OH 43210

**Author notes:** Corresponding author: Karen C. Dannemiller, Department of Civil, Environmental & Geodetic Engineering, Environmental Health Sciences, Sustainability Institute, Ohio State University, 470 Hitchcock Hall, 2070 Neil Ave, Columbus, OH 43210.

## Abstract

Respiratory viral illnesses are commonly spread in the indoor environment through multiple transmission routes, including droplets, aerosols, and direct/indirect contact. Indoors, resuspension of dust from flooring is a major source of human exposure. However, it is critical to determine viral persistence on dust and flooring to better characterize human exposure. The goal of this work is to determine viral viability on two carpet types (cut and looped) and house dust over time and after four different cleaning methods. MS2 and Phi6 bacteriophages were used to represent non-enveloped and enveloped viruses, respectively. These viral surrogates were placed in an artificial saliva solution and nebulized onto carpet or dust. Viability was measured at various time points (0, 1, 2, 3, 4, 24, and 48 hours) and after cleaning (vacuuming, hot water extraction with stain remover, steam, and a disinfection spray). Viability decay was modeled as first-order. MS2 bacteriophages showed slower viability decay rates in dust (−0.11 hr^-1^), cut carpet (−0.20 hr^-1^), and looped carpet (−0.09 hr^-1^) compared to Phi6 (−3.36 hr^-1^, -1.57 hr^-1^, and - 0.20 hr^-1^ respectively). The difference between phages was statistically significant in dust and cut carpet (p<0.05). Viral RNA demonstrated minimal degradation that in most cases was not statistically different from zero over the 48 hours measured (p>0.05). Viable viral concentrations were reduced to below the detection limit for steam and disinfection for both MS2 and Phi6 (p<0.05), while vacuuming and hot water extraction with stain remover showed no significant changes in concentration from uncleaned carpet (p>0.05). This study used viral surrogates and did not model risk of viral transmission via dust. Overall, these results demonstrate that MS2 and Phi6 bacteriophages can remain viable in carpet and dust for several hours to days, and cleaning techniques with heat and disinfectants may be more effective than standard vacuuming for viral removal. Future work should model risk from exposure via dust and flooring for various viruses such as influenza, SARS-CoV-2, and RSV.

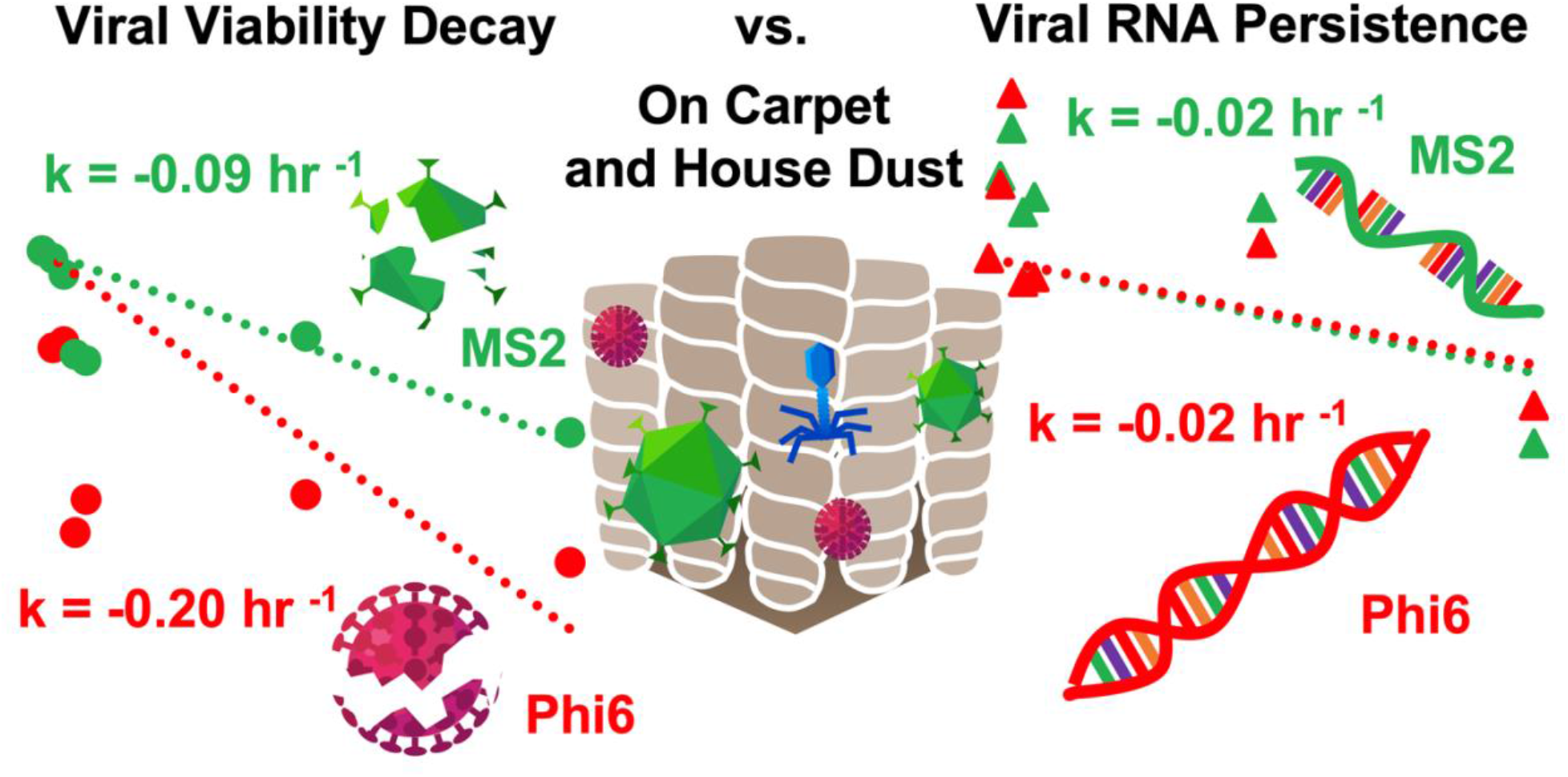

## Introduction

The novel severe acute respiratory syndrome coronavirus 2 (SARS-CoV-2) has resulted in more than 160 million cases and 3.3 million deaths worldwide [1] since reaching pandemic designation in March 2020. Transmission occurs predominantly in the indoor environment [2]. Spread occurs primarily through droplets and aerosols, though fomite transmission may also contribute at a lesser level [3–5] [6][7,8]. Dust represents an intermediate material that can be conceptualized as either a fomite or an aerosol. In fact, other respiratory viruses can be transmitted via particulate matter and have been conceptualized as ‘aerosolized fomites’ [9,10].

Viruses, including SARS-CoV-2, can persist on contaminated surfaces or materials [5]. In fact, SARS-CoV-2 remains viable on plastics and stainless steel, with a half-life on the order of hours [4]. A norovirus outbreak in school children followed a contamination event from an infected individual who vomited in the building on the previous day [11]. Typically, enveloped viruses (e.g., SARS-CoV-2, influenza) decay more rapidly on surfaces than non-enveloped viruses (e.g., norovirus, adenovirus) [12]. Environmental conditions such as relative humidity and the composition of carrier droplets also impact virus survival. Viruses typically remain viable the longest at low relative humidity levels, and viability may decrease as relative humidity increases or demonstrate a U-shaped pattern depending on droplet composition and presence of a viral envelope [13]. SARS-CoV-2 is also temperature sensitive [14].

RNA from SARS-CoV-2 and other viruses may be present at high levels in dust [15], and viruses on the floor are rapidly transported to hands and other surfaces [16]. SARS-CoV-2 has been detected on outdoor particulate matter [17]. Some evidence indicates that high particulate matter levels are associated with increased measles spread [18,19], although higher exposure to particulate matter could influence susceptibility separately. In fact, transmission of influenza via dust has been demonstrated in guinea pigs [9], and dust may contribute to spread of avian influenza [10]. However, viral persistence on indoor dust is not well understood.

The goal of this study is to determine persistence of two representative RNA viruses, Phi6 and MS2, on indoor dust and on carpet. Both viruses are bacteriophages and have widely been applied as surrogates for assessment of environmental fate of pathogenic viruses [20,21]. Phi6 (Φ6) has an enveloped capsid and MS2 is non-enveloped. In this study we assess viral viability over time and following four common cleaning measures: disinfection, vacuuming, steam cleaning, and hot water extraction with stain remover. Persistence was assessed both by culture (i.e., viability) and RT-qPCR (i.e., RNA detection) methods. Results of this work have important implications for understanding viral transmission in the indoor environment. This may also inform recommendations for cleaning practices following viral contamination.

## Methods

### Overview

The persistence of viable virus in house dust and residential carpet was observed in this study by using viral surrogates, MS2 and Phi6 bacteriophages. MS2 infects *Escherichia coli*, does not have an envelope, and has single-stranded, positive-sense RNA. Phi6 infects *Pseudomonas syringae*, has an enveloped capsid, and has double-stranded RNA.

To simulate viral deposition, the viral surrogates were placed in an artificial saliva mixture and nebulized onto carpet and dust samples. The virus was extracted from each sample using a wash and filtration step. We also evaluated carpet cleaning methods including vacuuming, steam cleaning, hot water extraction with stain remover, and disinfection to examine the effectiveness for inactivation or removal of the viral surrogates. For all samples, an RNA extraction and RT-qPCR analysis was performed to determine RNA quantity for each virus. Plaque assays were performed to determine viability.

### Carpet and Dust Samples

Carpet samples were supplied by a major manufacturer with no antimicrobial, stain resistance, or soil resistance coatings. The carpets were composed of polyethylene terephthalate (PET) carpet fibers and a synthetic jute backing. Two types of fiber construction processes were examined that included a cut pile (fiber length 10 mm) and a looped pile (fiber length 7.5 mm). Carpet samples consisted of a 5 cm x 5 cm square that contained a 3 cm x 1 cm cutout in the center that was used for viral viability and viral RNA assays to avoid edge effects. Triplicate carpet samples were used for each fiber construction type at each time point.

House dust was collected from a residential home vacuum bag in Ohio, USA. This dust was homogenized using a 300 µm sieve and confirmed to be negative for SARS-CoV-2 before use, using the IDT SARS-CoV-2 (2019-nCoV) CDC qPCR Probe Assay (Integrated DNA Technologies, Inc., Coralville, IA, USA) as described previously [15]. Dust was placed in sterilized glass dishes lined with tin foil that was previously baked (500°C for 12 hours). Each dish contained two 50 mg aliquots and a total of 3 dishes (6 dust piles) were used for each time point. One pile in each dish was used for viability assays and the other was used for RNA extraction and quantification. Carpet and dust samples were sterilized by autoclaving for 1 hour at 121°C and then placed in a 100°C oven overnight (∼12 hrs) to dry. Collection of dust for this study was approved by the Ohio State University Institutional Review Board (Study 2019B0457).

### Nebulization onto carpet and validation

Artificial saliva was created using a modified recipe [22](Table S1). Porcine gastric mucin was obtained from Sigma-Aldrich (Type II, M2378) [23]. We selected 10^8^ plaque forming units (PFU)/mL for both MS2 and Phi6 virus as a starting concentration to mimic the viral concentration of SARS-CoV-2 typically found in saliva, which ranged from 10^4^ - 10^8^ copies/mL one week after symptom onset [24]. Average viral concentrations in respiratory secretions and sputum range from 2.3×10^5^ virions/mL to 1.9×10^7^ virions/mL, depending on the time after symptom onset [25] [26], while peak viral load appeared 10 days after symptom onset [25]. This range of viral concentration in sputum was found to be similar in patients with both severe and mild SARS-CoV-2 symptoms [27].

A total of 4 mL of the saliva with both MS2 and Phi6 (10^8^ PFU/mL each) was nebulized onto the carpet or dust. Nebulization of viruses onto samples were performed following a modified protocol that increased the nebulizing and settling time to 15 minutes [28]. A total of three nebulization runs were conducted for each time point. Each nebulization deposited the viral saliva solution onto triplicate cut carpet, looped carpet, and dust samples. The triplicate samples for each material type were then placed into separate incubation jars.

### Incubation

After nebulization, samples were placed into a 3.8 L autoclaved glass jar for incubation. A salt solution (500 mL DI water, 268 g MgCl_2_) was made and placed in each jar to keep the equilibrium relative humidity (ERH) in the incubation chamber between 30-40%. Water activity of the salt solution was confirmed with an AquaLab 4TE (Meter Group, Pullman, WA). An Onset® HOBO® logger (Bourne, MA USA) was placed in the chamber to record temperature and ERH levels during the incubations. Actual ERH measurements varied depending on the sample with cut carpet, looped carpet, and house dust all having different peak ERH and a different amount of time that it took to reach equilibrium (Figure S1). Incubation time points that were sampled included a time 0 (immediately after nebulization) followed by 1, 2, 3, 4, 24, and 48 hours at which time the virus was extracted from each sample. All incubation jars were placed in a VWR incubator at 25°C.

### Carpet Cleaning

Several cleaning methods were evaluated for their effectiveness of inactivating Phi6 and MS2 bacteriophages. For these tests, cut carpet samples were nebulized with saliva/viral mixture as described above and cleaned immediately after nebulization (Time 0). Carpets were cleaned using vacuuming, hot water extraction with stain remover, steam, and application of a disinfectant solution. Each cleaning method was employed for 1 minute and then viruses were extracted from each carpet sample. A standard canister vacuum was used for vacuuming and a portable hot water extraction carpet cleaner was used for hot water extraction. The carpet cleaner was equipped with a 3-inch cleaning attachment tool with bristles for water extraction. A soap solution of 500 mL of water and 75 mL of a commercially available carpet stain remover was heated to 60°C then added to the cleaning tank. Using the cleaning attachment tool, 3 sprays of the cleaning solution were applied to the carpet samples. The solution was allowed to sit for 10 seconds and was then removed using the attachment tool. For steam cleaning, water was boiled to 100°C in a 200 mL glass beaker. The carpet was placed upside down on top of the beaker so that the steam made contact with the carpet (Temperature on carpet backing measured 80°C). For disinfection, a disinfectant spray was created with active ingredients sodium troclosene (NaDCC) and hypochlorous acid (HOCl). The solution was diluted to a 1076 ppm available chlorine using 10 tablets in 946 mL of DI water. One spray (∼2 mL) was applied to each carpet sample and allowed to sit for 1 minute before viral extraction. The disinfectant could not be removed prior to washing, so disinfection may have continued in subsequent steps. All cleaning methods were compared to carpets that were not cleaned but were nebulized with the same solution over two experimental trials.

### Viral extraction

For carpet samples, the pre-cut 3 cm x 1 cm rectangles were pushed out and placed in a 50 mL plastic tube with 8 mL of PBS to wash the virus from the material similar to previous extraction methods on fabric [29]. For all house dust samples, cups contained two separate 50 mg pre-weighed aliquots for each triplicate sample of which one was used for viability, and one was used for RNA analysis. The carpet/dust and PBS were mixed by hand then vortexed. A total of 4 mL of the wash was extracted and placed in a Amicon® Ultra-4 Ultracel®-50k (Merck Millipore Ltd.) filter tube and centrifuged for 7 minutes at 7000 rpm. The filtrate collected on the top of the filter was collected into a 1.5 mL tube and the volumes recorded for each sample. The process was repeated a second time using another 4 mL of PBS wash in order to collect enough for viability and RNA analyses.

For viability analysis, a dilution series of 9 was made for each sample using 100 µL of the collected sample and 900 µL of PBS. More details are below.

Phi6 and MS2 were extracted from samples of nebulized carpet utilizing the QIAamp DSP Viral RNA Mini Kit (Qiagen, Germantown, MD) and from samples of dust utilizing the RNeasy PowerMicrobiome Kit (Qiagen, Germantown, MD) respectively. 140 uL of the homogenous mixture of virus and PBS was placed in the lysis tube from the QIAamp DSP Viral RNA and the extraction protocol given in the kit was followed. To extract from the dust samples, 50 mg of nebulized dust was utilized following a modified RNeasy PowerMicrobiome Kit protocol using 10x the procedure recommended 2-mercaptoethanol and phenol chloroform based lysis [15].

### Viral propagation and enumeration

#### Phi6 bacteriophage

*Pseudomonas syringae* (Phi6 bacteriophage host) was grown from a frozen stock (supplied by Dr. Karen Kormuth at Bethany College) on 1.5% Luria-Bertani (LB) agar plates (20 g/L Difco™ Miller LB Broth, 10 g/L Bacto™ Agar) for 48 hours at 25°C. *P. syringae* was then transferred to an LB Broth (1 L DI water, 20 g Difco™ Miller LB Broth) one colony was used per 8 mL of LB Broth. This liquid culture was incubated for 16 hours at 25°C while shaking at 180 rpm. Phi6 bacteriophage (supplied by Dr. Karen Kormuth at Bethany College) was propagated using an enhanced MgCl_2_ solution (50 mL DI water, 50 mL LB, 1.25 mL of 1M MgCl_2_, 5 mL of *P. syringae* overnight culture, 20 µL of stock Phi6) incubated for 24 hours at 25°C. The enhanced solution was centrifuged for 30 minutes at 4000 rpm and filtered through a 0.22 µm filter. The high-titer Phi6 solution was made into a 40% glycerol solution and stored - 80°C until use. For enumeration of Phi6 bacteriophage, a 0.75% LBA (20 g/L Difco™ Miller LB Broth, 7.5 g/L Bacto™ Agar) was made and when cooled to 48°C was infused with *P. syringae* overnight culture (1 mL *P. syringae* per 10 mL of soft LBA). A total of 10 mL of the infused soft agar was pipetted into each culture plate. After the agar cooled, spot plating (Beck et al 2009) of each sample dilution was performed by using six 10 µL drops and was incubated for 24 hours at 25°C. Plaques were counted with a Darkfield Quebec® Colony Counter (Reichart, Inc. Depew, NY, USA) and PFU/mL was calculated. This value was converted to PFU per square centimeter of carpet and milligram of dust.

#### MS2 Bacteriophage

*Escherichia coli* F_amp_ (MS2 bacteriophage host) cultures were incubated in LB liquid media from a frozen stock (supplied by Dr. Karl Linden and Dr. Ben Ma at University of Colorado Boulder) on a shaker table at 180 rpm and 36°C for 16 hours. After incubation, 1.467 mL of this overnight culture was transferred to 200 mL of LB and incubated at 225 rpm for an additional 2.5 hours. For propagation of MS2 bacteriophage, 10 mL of this culture was transferred to a new flask where 1.267 mL of 1M MgCl_2_ and 633 µL of frozen MS2 stock (supplied by Dr. Karl Linden and Dr. Ben Ma at University of Colorado Boulder) were added. The solution was gently mixed and allowed to sit for 25 minutes before resuming incubation at 36°C at 185 rpm for another 2.5 hours. After propagation, cultures were centrifuged at 7000 rpm and 10°C for 15 minutes. The supernatant was aliquoted into 1 mL stocks and stored at -80°C until use. For MS2 enumeration, a 0.75% LBA was made and when cooled *E. coli* F_amp_ from the 2.5-hour incubation was added (200 µL *E. coli* F_amp_ per 10 mL of 0.75% LBA). The samples were spotted on to the plates using the same method as the Phi6 bacteriophage and were incubated at 36°C for 24 hours at which time PFU/mL was counted and calculated in the same manner as Phi6 bacteriophage.

### RT-qPCR

In preparation for cDNA synthesis, heat shock treatments were performed to denature the dsRNA segments in the Phi6 genome. For heat treatment, 5 uL of sample was held at 100°C for 5 min followed by 5 min on ice as recommended by Gendron (2010). A heat shock treatment was not used for MS2 genomes. cDNA was reverse transcribed from RNA samples using the iScript cDNA Synthesis Kit (Biorad, Hercules, CA) according to the recommended reaction protocol on the ProFlex PCR System (Applied Biosystems, Forest City, CA). The cDNA was stored at -80°C. MS2 and Phi6 specific primers and probes were used to determine RNA concentrations. The MS2 forward primer (5’-GTCCATACCTTAGATGCGTTAGC-3’), reverse primer (5’-CCGTTAGCGAAGTTGCTTGG-3’), and probe (5’-/56-FAM/ACGTCGCCAGTTCCGCCATTGTCG/3BH) and the Phi6 forward primer (5’-TGGCGGCGGTCAAGAGC-3’), reverse primer (5’-GGATGATTCTCCAGAAGCTGCTG-3’) and probe (5’-/5FAM/CGGTCGTCGCAGGTCTGACACTCGC/3BH) were used in the PCR reactions [30]. The PCR final reaction mixture contained 1X TaqMan® master mix (Applied Biosystems™), 1 µM of forward and reverse primers, 150 nM of MS2 probe or 300nM of Phi6 probe, and 2 µl of cDNA template and the volume was adjusted with sterile water to 25 µL [30].

Quantitative polymerase chain reaction (qPCR) was completed on a QuantStudio™ 6 Flex System (Applied Biosystems™) with samples prepared on a 384-well (0.2 mL) plate in triplicate. Amplification protocol consisted of 50°C for 2 min 95°C for 10 min followed by 40 cycles of 94°C for 15 sec and 60°C for 1 min.

### Statistical analysis

We evaluated decay of viral viability and RNA persistence over time on carpet and dust as well as the reduction in viral viability after use cleaning techniques. The decay curves from all experiments were fit to a first-order decay model. The first order decay rate constant, k (h^-1^), can be calculated as the slope of the line ln (C_t_/C_0_) versus time where C_t_ is the concentration of the virus at any time t and C_0_ is the initial concentration of the virus at time zero. The mean of triplicate measurements was calculated at each time point and used in the simple linear regression analysis to calculate the first order decay rate constant. All decay rate constants, regression coefficients and confidence intervals were calculated using GraphPad PRISM ver. 9 (GraphPad, San Diego, CA) fixing the y-intercept to zero. Confidence bands represent the boundary for all possible lines and were determined and plotted using GraphPad PRISM ver. 9. Statistical correlation between the decay of MS2 and Phi6 bacteriophages in the two different types of carpet (cut and loop) and dust was determined using Analysis of Covariance (ANCOVA). This analysis was also performed to determine correlation between the decay of the two different types of bacteriophages (MS2 and Phi6) in dust and artificial saliva. Phi6 bacteriophage viability decay in saliva alone was not statistically different from zero whereas MS2 decay in saliva had a small growth rate that was statistically different from zero but much less than the rates for decay on carpet/dust (p=0.001 for MS2, p=0.098 for Phi6) (Figure S2, Table S2). Comparison between untreated cut fiber carpet and carpet cleaned by vacuum, steam, hot water extraction, and disinfection was done for Phi6 and MS2 bacteriophage using Kruskal-Wallis test followed by Dunn’s multiple comparisons test. Values below detection limit were assumed to be half detection limit for Kruskal-Wallis test.

## Results

### Viral decay over time

In house dust, Phi6 had a faster viability decay rate (−3.36 hr^-1^) compared to MS2 bacteriophage (−0.11 hr^-1^) (Figure 1, Table 1), and the difference was statistically significant (ANCOVA, p=0.0001). Phi6 had faster viability decay than MS2 in both looped and cut carpet type, and the difference was statistically significant for cut carpet (ANCOVA p=0.0018 for cut and p=0.20 for looped).

**Table 1:**
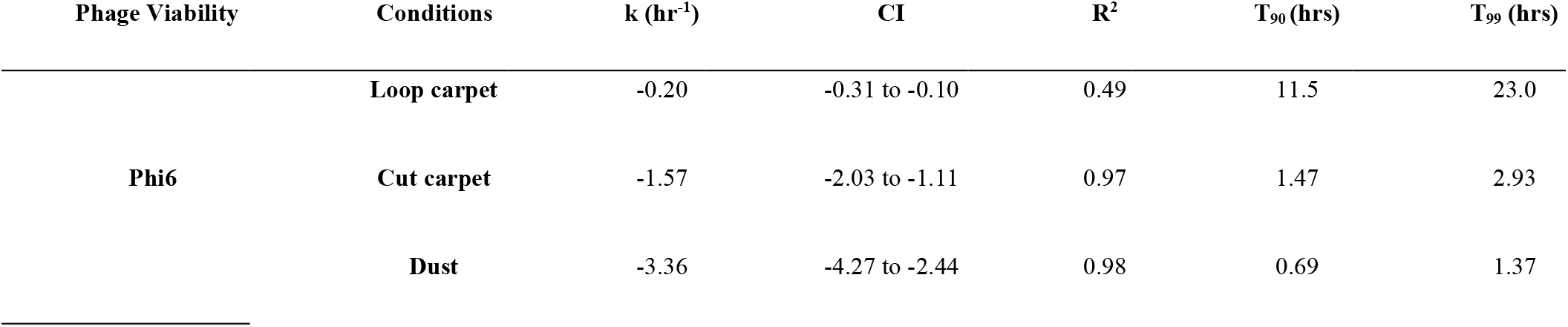
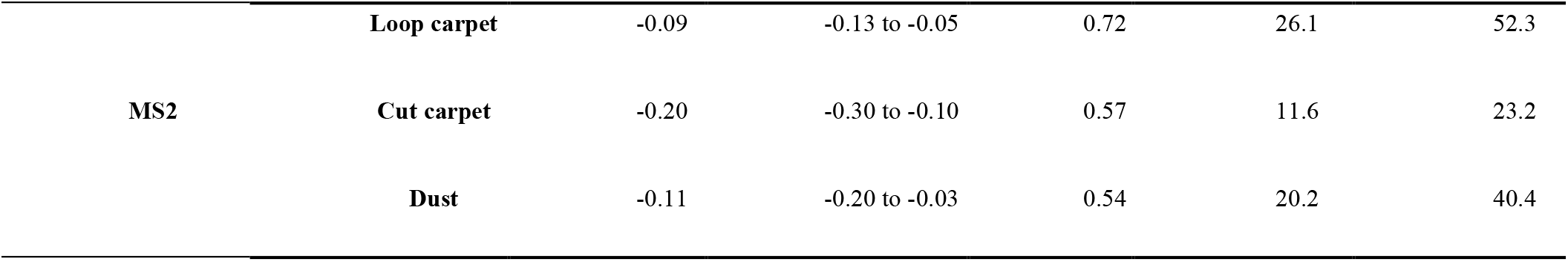
Viability first-order decay rate constants for Phi6 and MS2 bacteriophage in house dust, cut and loop carpet types.

**Figure 1:**
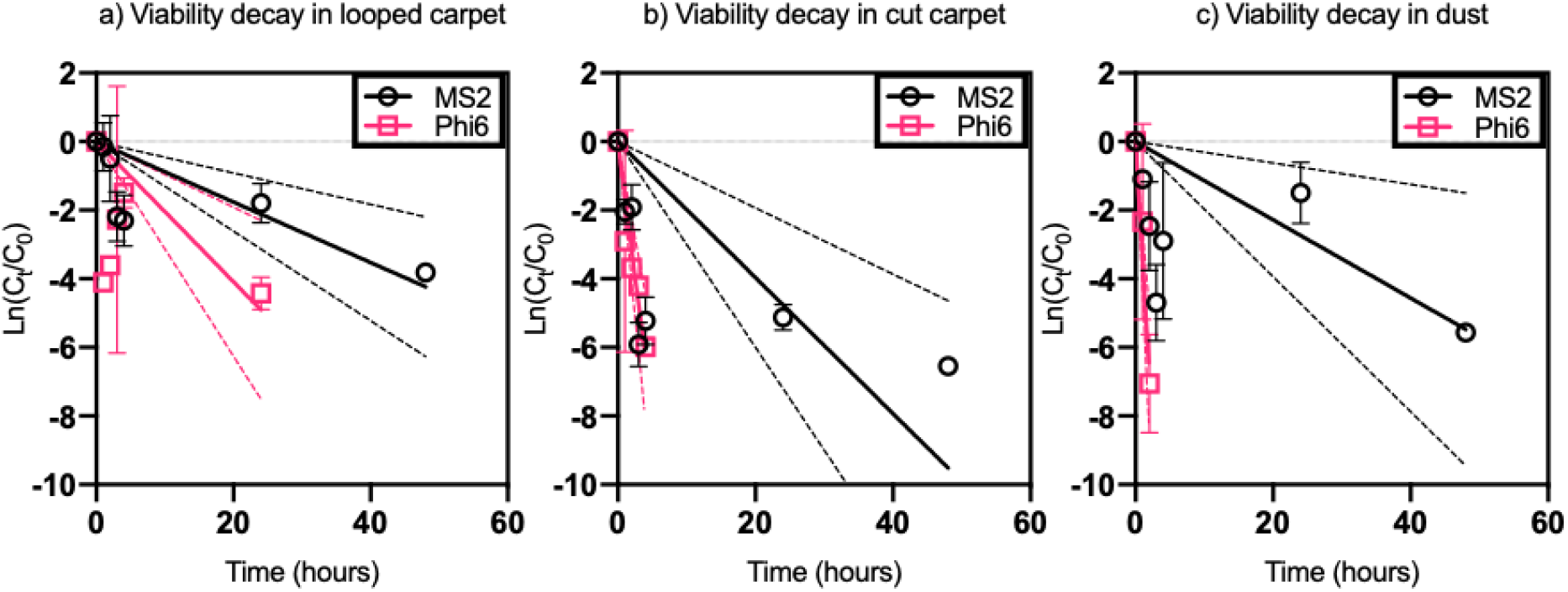
Decay of MS2 and Phi6 bacteriophage viable virus in loop (a) and cut (b) carpet types and dust (c). Each data point represents the average of experimental triplicate measurements from each sample type (cut carpet, looped carpet, and dust) at each time point. Each point represents mean sample concentration and error bars represent the standard deviation for each sample. Viable Phi6 was detectable for 24 hours for looped carpet, 4 hours for cut carpet, and 2 hours in dust. Viable MS2 was detectable for the full duration of the experiment (48 hours) in all conditions. Dashed lines represent 95% confidence bands for regression lines. Detection limits for carpet, house dust, and plaque assays were 6 PFU/cm^2^, 0.52 PFU/mgt, and 16.7 PFU/mL respectively.

In the artificial saliva, MS2 had a slight positive slope and the viability decay rate for Phi6 was not statistically different from zero (Figure S2, Table S2). The viability decay rates were statistically different from each other (ANCOVA, p=0.0002).

For both Phi6 and MS2 bacteriophages, viability decay occurred faster in cut carpet fibers (−1.57 and -0.20 hr^-1^) compared to looped carpet fibers (−0.20 and -0.09 hr^-1^), and the difference was statistically significant for Phi6 (ANCOVA p=0.0011 for Phi6, p=0.35 for MS2). Additional data including viability values and viable concentrations plots can be found in the supporting information (Figures S3-S5, Tables S3-S5).

Decay of RNA was much slower compared to decay of viability in both carpet types and in dust for both Phi6 and MS2 (Figure 2, Table 2). Decay of Phi6 RNA was not significantly different from decay of MS2 RNA in looped and cut carpet but was significantly different in dust (ANCOVA p=0.67 for loop, p=0.91 for cut and p=0.042 for dust). MS2 RNA decay was also not statistically different from zero in dust, looped and cut carpet (ANCOVA p=0.11 for loop, p=0.12 for cut and p=0.31 for dust). Phi6 RNA decay in dust and cut carpet was statistically different from zero whereas decay in looped carpet was not statistically different from zero (ANCOVA p=0.01 for dust, p=0.03 for cut and p=0.06 for looped). There was also no statistical difference in Phi6 RNA decay between dust, looped and cut carpet types (ANCOVA p=0.07 for looped/cut, p=0.19 dust/cut and p=0.57 dust/loop). Additional data including raw qPCR values and RNA concentrations plots can be found in the supporting information (Figures S6-S8, Tables S6-S8).

**Table 2:**
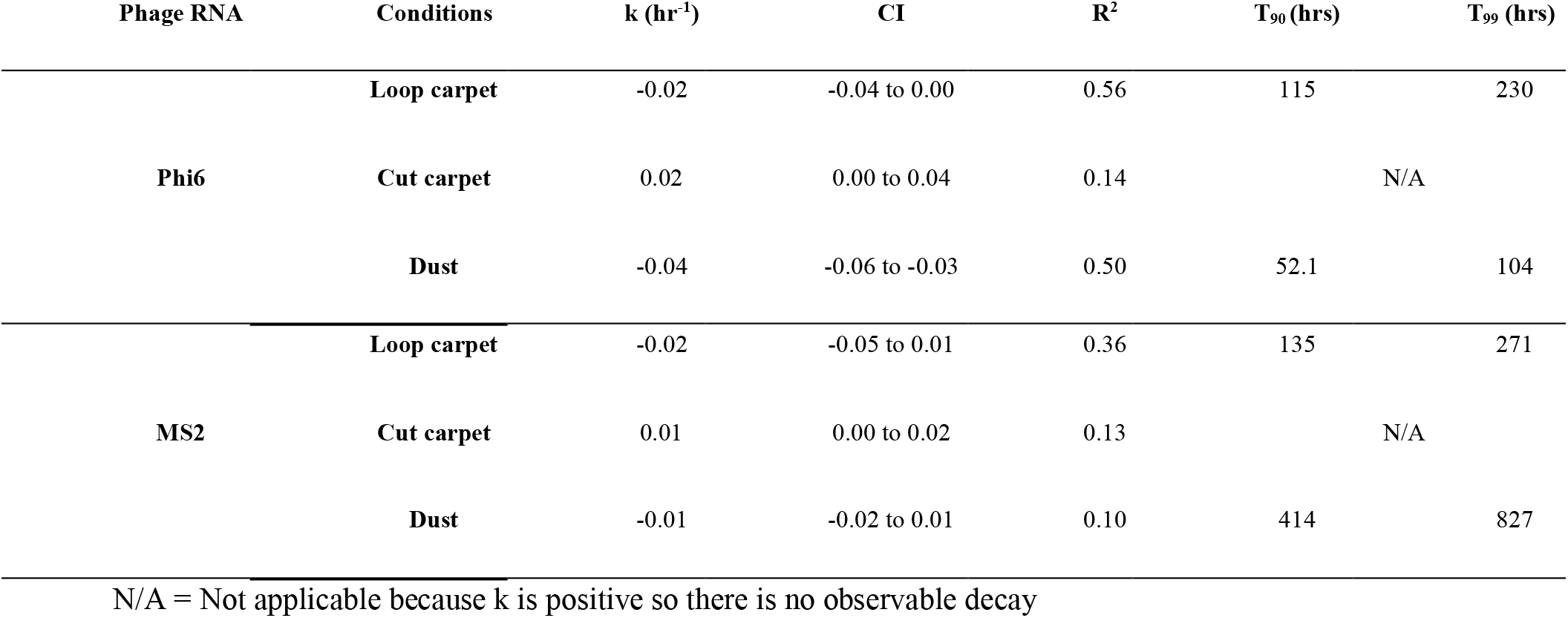
RNA first-order decay rate constants for Phi6 and MS2 bacteriophage in house dust, cut and loop carpet types.

**Figure 2:**
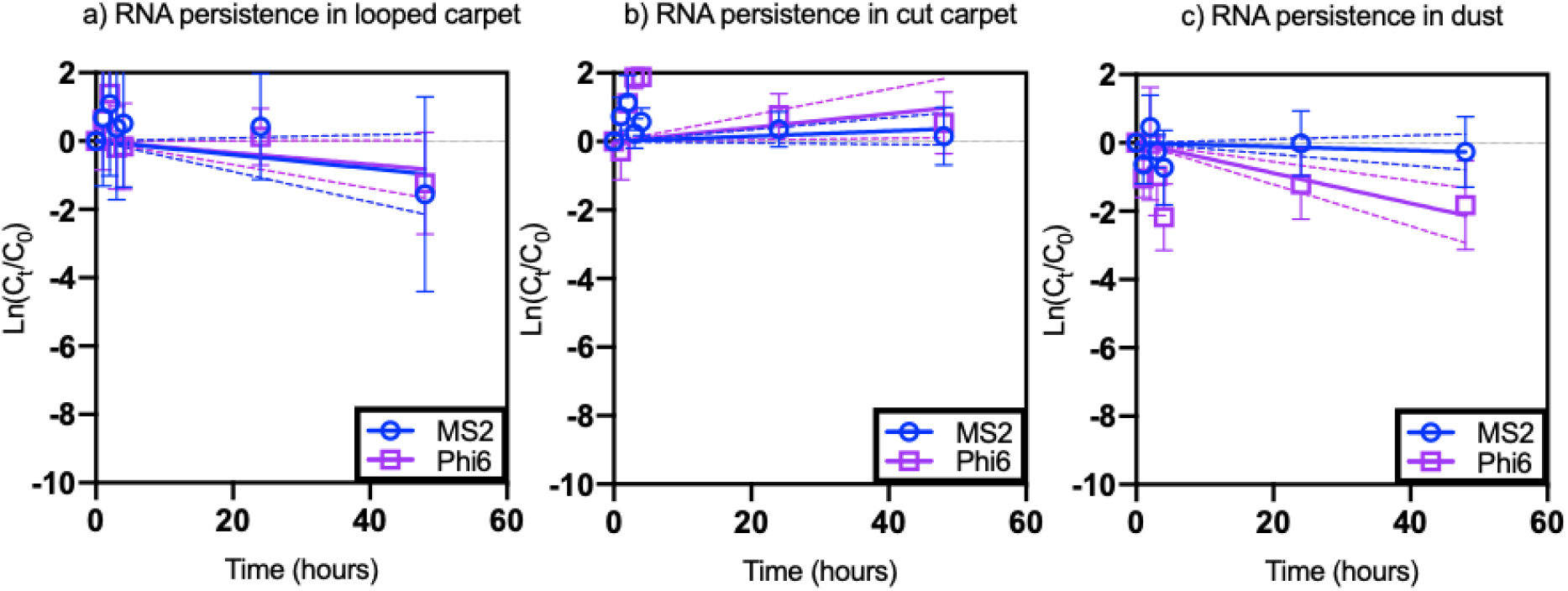
Decay of MS2 and Phi6 bacteriophage RNA in loop (a) and cut (b) carpet types and dust (c). Each data point represents the average of triplicate measurements. Each point represents mean sample concentration and error bars represent the standard deviation for each sample. MS2 and Phi6 RNA were detectable for the full duration of experiment (48 hours) in all conditions. Dashed lines represent 95% confidence bands for regression lines. Detection limits for carpet, house dust, and plaque assays were 6 PFU/cm^2^, 0.52 PFU/mgt, and 16.7 PFU/mL respectively.

### Removal after cleaning treatments

We compared concentrations of viable MS2 and Phi6 bacteriophage as well as concentrations of RNA from MS2 and Phi6 bacteriophage on untreated (no cleaning), vacuumed, steam treated, disinfected, and hot water extracted carpet (Figure 3). Viability concentrations were below the detection limit for cut carpet cleaned with steam and disinfectant for both Phi6 and MS2, and concentrations of Phi6 (Kruskal-Wallis, p=0.014 for steam and 0.030 for disinfectant) and MS2 (Kruskal-Wallis, p=0.015 for steam and p=0.028 for disinfectant) bacteriophage were statistically different on these treated carpets compared to untreated carpet. Hot water extraction with stain remover and vacuuming left measurable viable MS2 and Phi6 bacteriophage on the carpet fibers, and concentrations of MS2 (Kruskal-Wallis, p=0.27 for hot water extraction and p=0.095 for vacuuming) and Phi6 (Kruskal-Wallis, p=0.28 for hot water extraction and p=0.62 for vacuuming) were not statistically different on these treated carpets compared to untreated carpet (Figure 3A).

**Figure 3:**
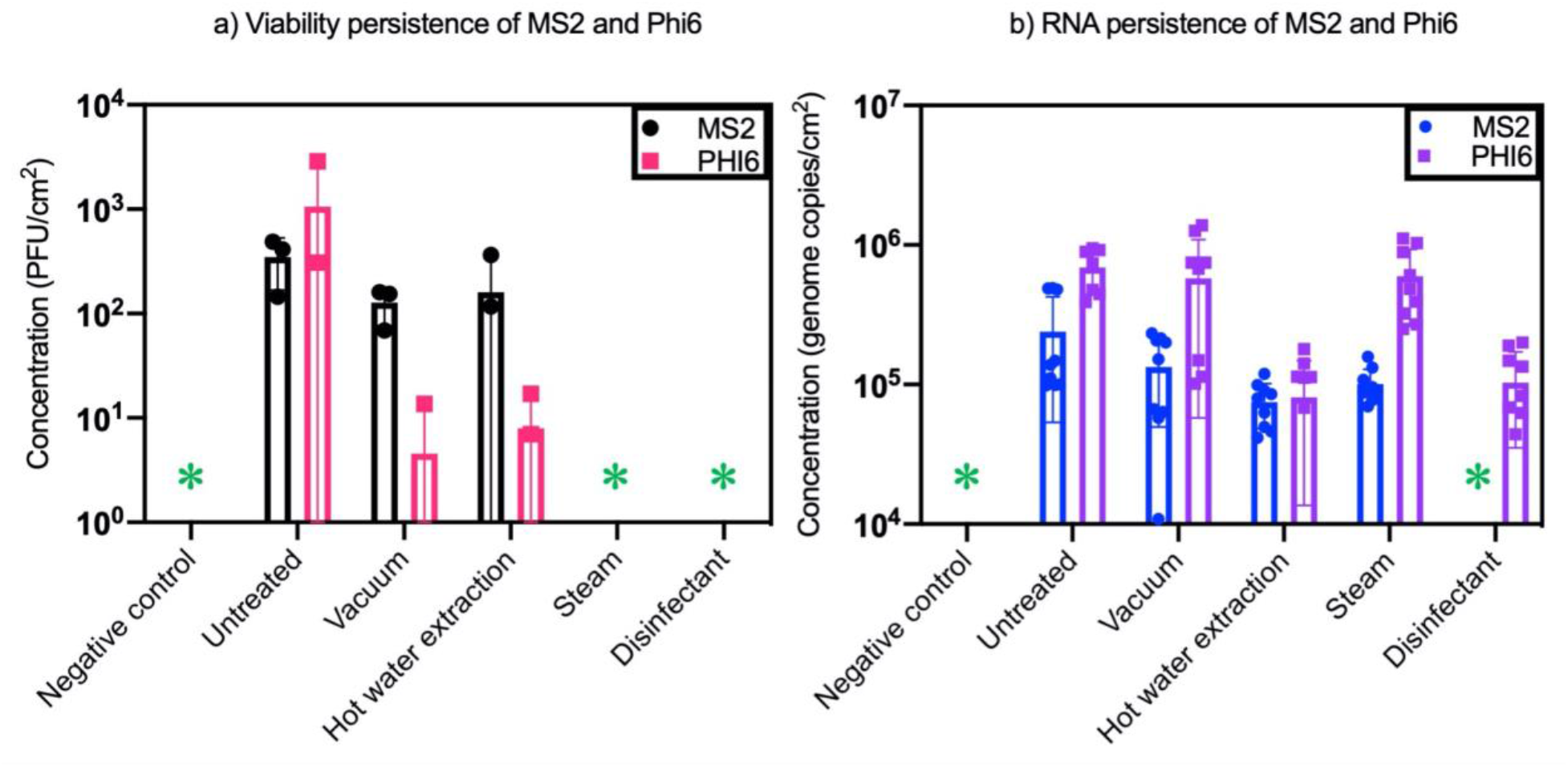
Concentrations of viable MS2 (black) and PHI6 bacteriophage (pink) on untreated (MS2 detected 3/3, PHI6 detected 3/3), vacuumed (MS2 detected 3/3, PHI6 detected 1/3), hot water extraction treated (MS2 detected 2/3, PHI6 detected 2/3), steam treated (MS2 detected 0/3, PHI6 detected 0/3) and disinfection treated (MS2 detected 0/3, PHI6 detected 0/3) carpets (a). Concentrations of RNA from MS2 (blue) and PHI6 (purple) bacteriophage on untreated, vacuumed, hot water extraction, steam treated, and disinfection treated carpets (b). Points represent concentration in each sample and error bars represent standard deviation. ^*^Represents values below the detection limit. Detection limit for carpets is 6 PFU/cm^2^.

Concentration of RNA in the untreated samples was not statistically different compared to vacuum cleaned or steam cleaned for Phi6 (Kruskal-Wallis p=0.29 for vacuum and p=0.75 for steam). Concentrations of RNA from carpets cleaned with hot water extraction and disinfectant were statistically lower than the untreated samples for Phi6 (Kruskal-Wallis p=0.0007 for hot water extraction and p=0.0023 for disinfectant). Utilizing a vacuum or steam cleaner did not influence the RNA concentration of MS2 when compared to the untreated samples (Kruskal-Wallis p=0.28 for vacuum and p=0.15 for steam). However, RNA concentrations of MS2 were statistically lower when utilizing hot water extraction or a disinfectant (Kruskal-Wallis p=0.016 for hot water extraction and p=0.0001 for disinfectant). Additional raw data values for cleaning method viral viability and RNA are found in Tables S9-S10.

## Discussion

Carpet and dust are potentially important reservoirs for microbial exposure to humans in the built environment because they serve as both a sink and a source for bacteria, fungi, and viruses [31]. Our work demonstrates that viruses can remain infective in dust for hours to days, and that the presence of a viral envelope may be an important factor in determining persistence time. Additionally, cleaning methods have a range of removal efficiencies, with methods that employ heat or disinfectants being more effective than vacuuming. RNA persisted on carpet and dust longer than viable viruses.

### Role of flooring in viral resuspension

Transmission of respiratory viruses occurs predominantly in the indoor environment, and it is critical to understand viral viability on flooring and dust because these are an important source of human exposure. Exposure to virus in flooring will be influenced predominantly by two factors: 1) presence of the virus, influenced by deposition and viability decay, and 2) resuspension into the breathing zone. Regarding the former, nucleotides from SARS-CoV-2 and other viruses, like influenza, are detectable in dust and air samples, but often viability is not measured [15,32–35]. In our study, PET carpet fibers with a synthetic jute backing material were utilized and the fiber construction varied between looped pile and cut pile. Both viruses were viable longer on the loop carpet compared to the cut carpet.

Resuspension of dust due to walking is an important contributor to human exposure, particularly in carpeted areas [36]. The resuspension of dust in flooring however is also likely impacted by type of flooring material such as looped carpet, cut pile carpet and hard flooring [37,38]. A recent study posited that a possible source of aerosolized SARS-CoV-2 is due to the resuspension of floor dust from walking in areas with confirmed positive patients [39]. Dust particles contaminated with influenza virus may be resuspended and serve as aerosolized fomites of viable influenza virus [9,40]. In fact, the concentration of the resuspended influenza virus was 40% higher at one meter than two meters, such that shorter people and children may be more likely to come in contact with these particles [40]. Understanding the resuspension of SARS-CoV-2 and other viruses in carpet will be important in measuring exposure route, and resuspension models are needed to determine the rate of resuspension of SARS-COV-2 in varying carpet types.

### Viral envelope may impact persistence on flooring and dust

The presence of carpets has been linked to viral infections, especially for nonenveloped viruses. In one instance, carpet fibers still contained Norwalk-like viruses 13 days after the last infection [41]. Outbreaks of the nonenveloped human norovirus have also been linked to viral transfer of surfaces such as cotton and polyester [42]. In one study, human norovirus surrogates were determined to survive for ∼15 days on carpet fibers depending on relative humidity condition and fiber type [43]. Non-enveloped viruses, such as rotavirus and poliovirus, persist for up to two months on surfaces [44,45], while enveloped respiratory viruses, such as H1N1, human coronaviruses and SARS-CoV, persist for several hours to days [46][47].

In our study, the non-enveloped viral surrogate MS2 persisted as viable in carpet longer than enveloped Phi6. As SARS-CoV-2 is an enveloped virus, it may persist in carpet more similarly to that of Phi6, and this can be evaluated in future studies. We used a first-order equation to model viral viability decay. Model fit may have been influenced by variability inherent in sample processing, which may be due to differences in nebulization, carpet wash steps, and viral plaque assays in general. However, the trend was clear that Phi6 viability decays more quickly than MS2. RNA from both bacteriophages persisted longer than viability.

In all cases the type of material does impact the persistence of the virus. Here, we saw different viability decay rates based on carpet fiber type and on carpet compared to dust. SARS-CoV-1 survives on metal, wool, paper, glass, and plastic for 4 - 5 days [48–50]. In other cases, on non-porous surfaces such as glass, stainless steel and vinyl, infectious SARS-CoV-2 was detected after 28 days; however, it was not detected on the porous material cotton cloth after 14 days at 20°C potentially due to an immediate absorption effect [14]. Persistence of SARS-CoV-2 on plastic (e.g., PET, which is the same material of our carpet) lasts for up to 72 hours and in one study had the longest viability of materials tested [4].

### Relative humidity and virus viability

Airborne viruses are sensitive to humidity conditions. Viruses remain viable the longest at low relative humidity, and viability may decrease as relative humidity increases or demonstrate a U-shaped pattern (high-low-high) depending on droplet composition and presence of a viral envelope [13]. SARS-CoV-2 is most stable at low relative humidity (<40%) and stability decreases as relative humidity is increased between 50 - 90% [51]. Virus viability rapidly decays at high temperatures (38°C) and high relative humidity (>95%) [49]. Other studies have determined that saturated humidity conditions may facilitate the spread of the SARS-CoV-2 via cluster spread [52]. At elevated humidity conditions between 65% - 100% aerosolized SARS-CoV-2 may be viable for up to 3 hours [4]. In this study the relative humidity was maintained ∼30% to mimic realistic home environment conditions, however due to incubation limitations this humidity level varied between 40-90% early in the incubation depending on the sample (Figure S1). The variation between carpet fiber structures could change the moisture retention properties and thus impact the decay rates for MS2 and Phi6 bacteriophages. However, this variation in carpet fiber construction is realistic in real-world building environments and should be investigated further to examine its effect on viral-saliva droplet deposition and evaporation. Elevated humidity conditions in carpet dust are known to influence fungal growth [53,54] and this humidity may influence the viability and stability of viruses in carpet dust. Continued work is needed to fully understand how elevated humidity in carpet may impact the viability as well as the stability of SARS-CoV-2 and other viruses in dust.

### Differential viral removal efficacy of cleaning methods

Inactivation or removal of viruses from house dust or residential carpets is an important tool that can be used to reduce viral transmission in indoor environments. Vacuuming is a common housekeeping routine used to remove accumulated soils from carpet. This method reduced viable viruses on our carpet samples, when compared to carpet samples that were not cleaned, but the viable virus was still detectable in the carpet afterward. However, vacuuming, and hot water extraction removed viable Phi6 (enveloped) more effectively compared to MS2 (non-enveloped). Vacuuming could also resuspend viral particles into the air [55]. Similarly, hot water extraction is often used to maximize physical removal of soils in carpet and is readily available for use in homes. This cleaning method reduced viable virus on carpet samples, but viruses were still detected after cleaning and the difference was not statistically significant. Applying steam to carpet samples reduced virus viability to below detectable limits but is not necessarily realistic for in-home decontamination of carpets. The application of a disinfectant to the carpet samples also reduced viruses to below detectable limits, although disinfectant may have continued to contact the virus during the wash step of our experiment. Disinfectants are an easy and realistic method building occupants may use to inactivate viruses on carpet following viral illness. These factors are likely to be most important in commercial/public, high-risk areas such as hospitals, and may only be relevant in residential settings under unique circumstances.

### Limitations

All carpet and dust samples were autoclaved before nebulization of viruses. However, due to the complexity of these samples it is likely that bacterial and fungal quantities were substantially reduced but not completely sterilized [56] A salt solution was used to attempt to maintain ERH in the incubation chambers at 30-40%. However, each sample type (dust, cut, and looped carpet) showed different absorption of nebulized saliva, which affected the peak ERH and the time in which it took to lower to under 50% ERH. Cut carpet peaked close to 90%, looped carpet approximately 80%, and dust at 60% ERH while taking 12, 6, and 3 hours respectively to reach 50% ERH. These differences in ERH and duration may have affected the decay rates of each viral surrogate in this study, although carpet structure might also reflect moisture retention in carpets in buildings. This may have also resulted in suboptimal first-order model fit, in addition to other factors that contributed to variation including variability in nebulization, carpet wash steps, and viral plaque assay limitations. Other models, such as biphasic decay, require more data points for accurate curve fit and could be evaluated in future studies. During aerosolization and deposition of the viral surrogates onto these samples, phage aggregates may have formed. This could create an uneven distribution among samples leading to an underestimation of the viable virus observed on carpet and in dust [30]. Different lab personnel performed the cleaning methods and time series viability testing, therefore user differences in the nebulization, carpet/dust extraction, and plaque assay protocols may have been introduced. For the disinfection cleaning method, we were unable to determine if viral viability was lost from contact on the carpet or while it was mixed during the viral wash extraction protocol.

This study used bacteriophage viral surrogates, which are different from human pathogenic viruses. Bacteriophages are considered good surrogates to study airborne viruses. These viruses can be produced in large quantities, pose little hazard for laboratory workers, and do not require specialized containment protocols [21]. The bacterial viruses MS2 and Phi6 were chosen for this work because of their similarities with known pathogenic viruses, including SARS-CoV-2 [20,21,57,58]. However, these are different viruses, and real pathogens may behave differently.

## Implications

These results suggest that viable viruses can persist on dust and carpet for hours to days depending on viral structure and environmental conditions, but more information is needed to understand risk. RNA persists longer than viable viruses and may be useful for surveillance methods [15]. This current study was based on bacteriophages, and additional research is needed using human viruses in the appropriate biosafety facilities to confirm the results, followed by risk modeling. Additionally, many enveloped viruses (including SARS-CoV-2 and others) may only remain infectious in dust for a very brief period of time (on the order of hours post-deposition). Thus, transmission may only be possible shortly after deposition, when expelled respiratory aerosols may also continue to be a transmission risk in the same space. Cleaning of such spaces could be delayed by several hours to reduce infection risk to maintenance staff. Non-enveloped viruses (norovirus and others) may be more easily transmitted via flooring over longer periods of time, and appropriate cleaning techniques using heat and/or disinfectants may be more critical to reduce infection risk. Unfortunately, in many cases the virus causing infection may not have been identified prior to the need for an environment to be cleaned, so it may be prudent to both wait some time if the room can be vacated and then employ heat-based or disinfection-based methods, when possible, to clean contaminated flooring.

Ultimately, future research can improve our understanding of dust as a potential transmission route for viral infection. Risk modeling should follow this analysis. This may have important implications for reducing viral spread in the general population, and also for custodial and cleaning staff who may be working closely to clean contaminated flooring. A more nuanced recognition of this potential exposure pathway can help contribute to the fight against viral illnesses such as influenza and COVID-19.

## Acknowledgements

We appreciate Dr. Karl Linden and Dr. Ben Ma at the University of Colorado Boulder for sharing MS2 bacteriophage and *E. coli* F_amp_. We also thank Dr. Karen Kormuth from Bethany College for sharing Phi6 bacteriophage and *Pseudomonas syringae*, as well as her culturing expertise related to these isolates. We thank the carpet manufacturer for the donation of carpet samples. This work was funded through faculty startup funds at The Ohio State University. We also want to acknowledge grant 1942501 from NSF, grant G-2018-1124 from the Alfred P. Sloan Foundation, and grant 80NSSC19K0429 from NASA, which allowed us to develop the expertise to conduct this analysis. This manuscript represents the views of the authors and has not been reviewed by funding agencies.

