## Supporting Information for "Viability of MS2 and Phi6 Bacteriophages on Carpet and Dust"

**Figure S5:** MS2 and Phi6 bacteriophage viability concentrations in house dust. Error bars represent 95% confidence intervals. "*" indicates below detection limits (0.52 PFU/mg dust). ... 5

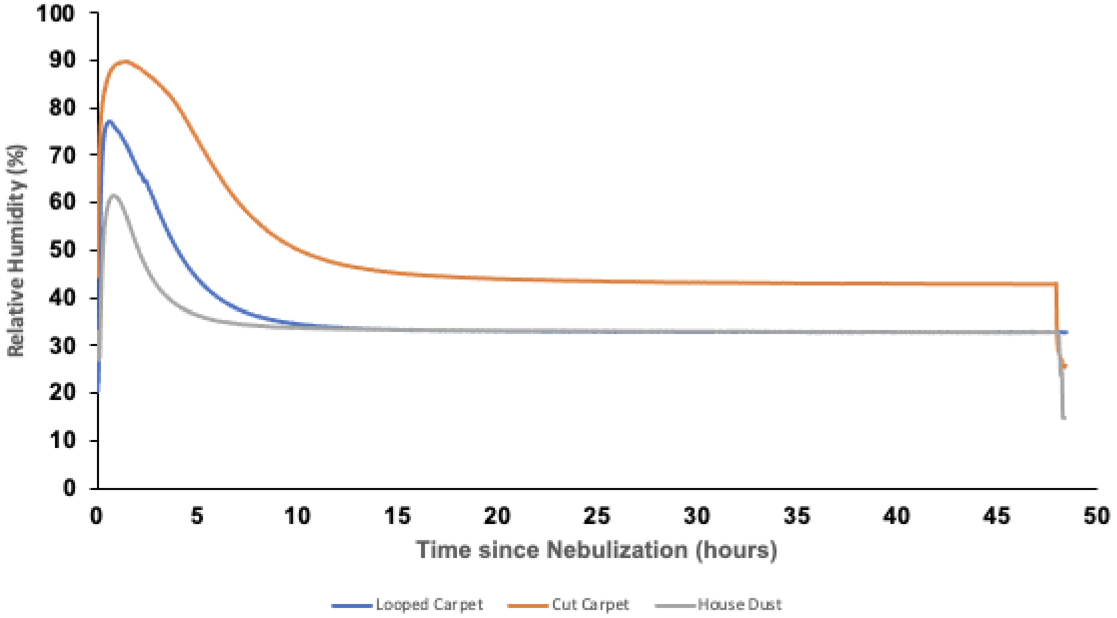

**Figure S1:** Equilibrium relative humidity measurements as recorded by HOBO® logger in incubation chambers for house dust (gray), cut carpet fibers (orange), and looped carpet fibers (blue). Each curve is from one chamber incubated for 48 hours and is generally representative of other incubation chambers at the shorter time periods examined in this study.

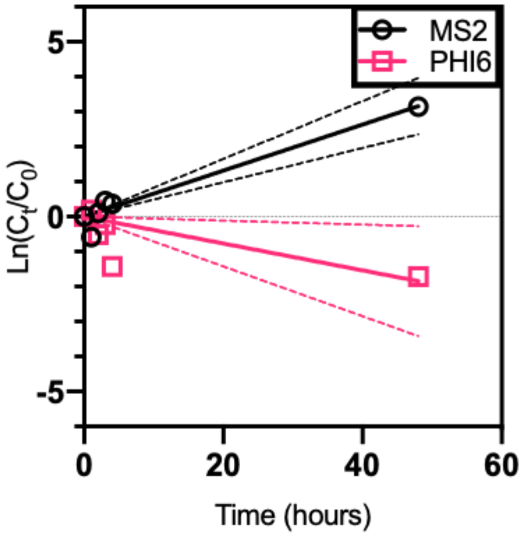

**Figure S2:** Viability decay of MS2 and Phi6 bacteriophages in artificial saliva. Each data point represents one measurement taken at each time point for each bacteriophage. Dashed lines represent 95% confidence bands for the regression lines.

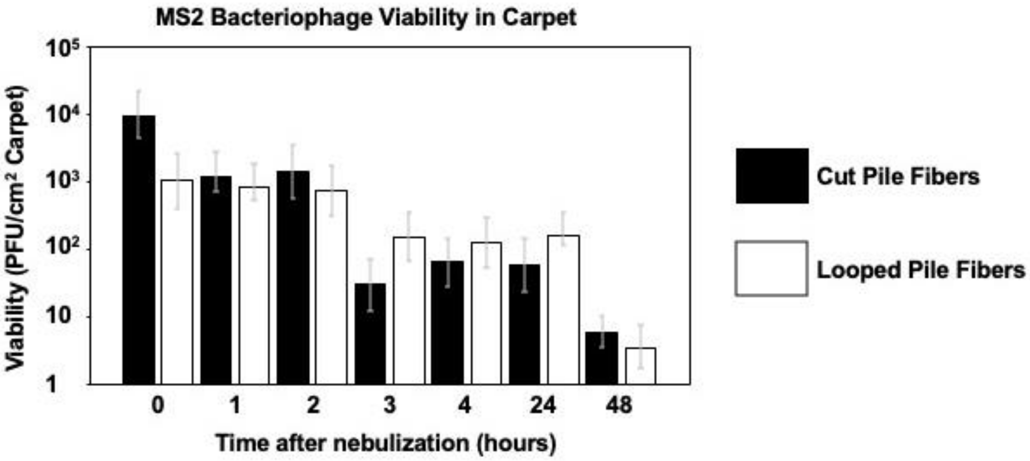

**Figure S3:** MS2 bacteriophage viability concentrations in cut and looped carpet fibers. Error bars represent 95% confidence intervals.

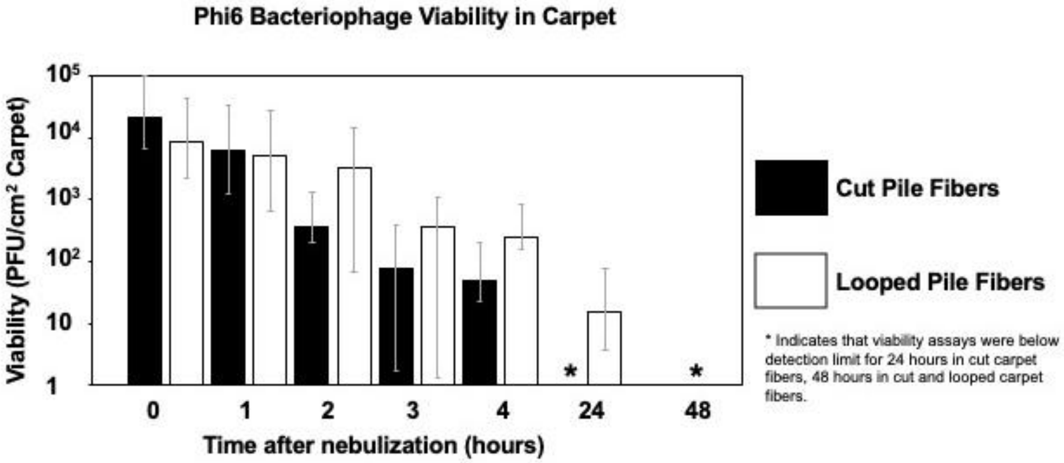

**Figure S4:** Phi6 bacteriophage viability concentrations in cut and looped carpet fibers. Error bars represent 95% confidence intervals. "*" indicates below detection limits (6 PFU/$cm^2$ carpet).

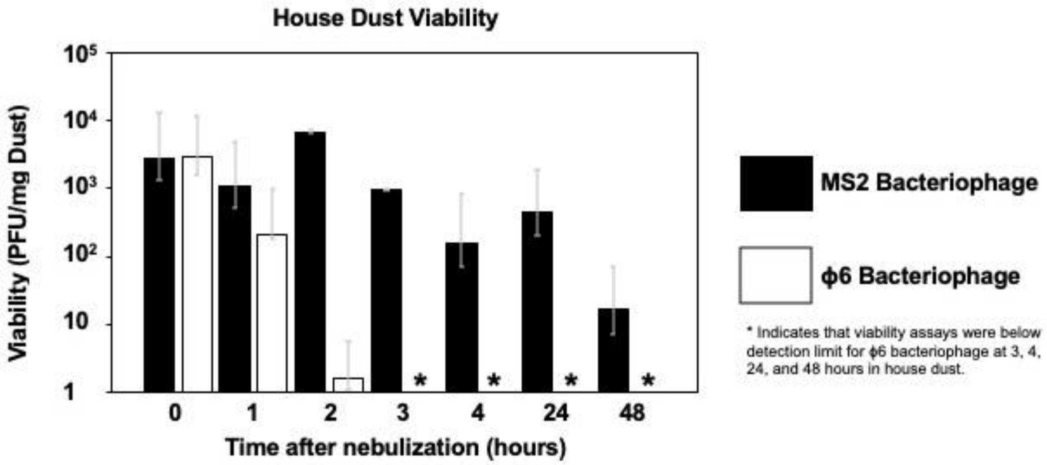

**Figure S5:** MS2 and Phi6 bacteriophage viability concentrations in house dust. Error bars represent 95% confidence intervals. "*" indicates below detection limits (0.52 PFU/mg dust).

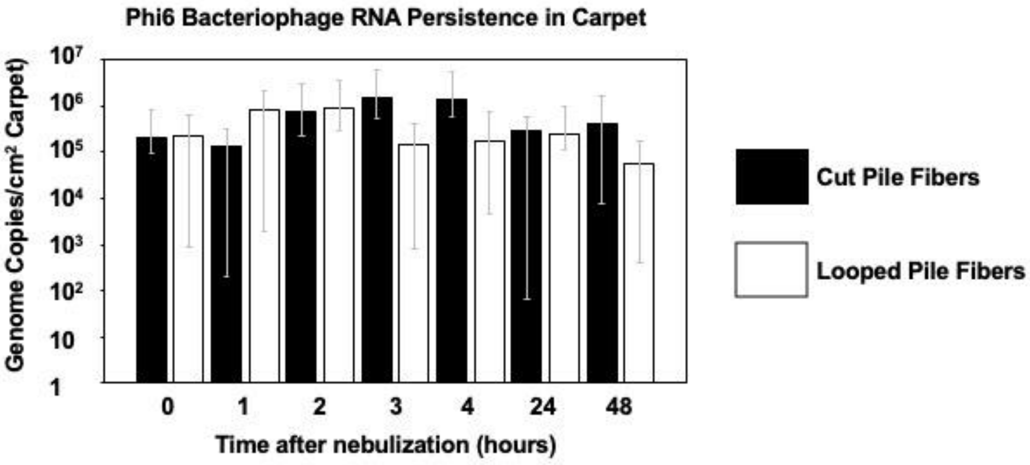

**Figure S6:** Phi6 bacteriophage RNA concentrations in cut and looped carpet fibers. Error bars represent 95% confidence intervals.

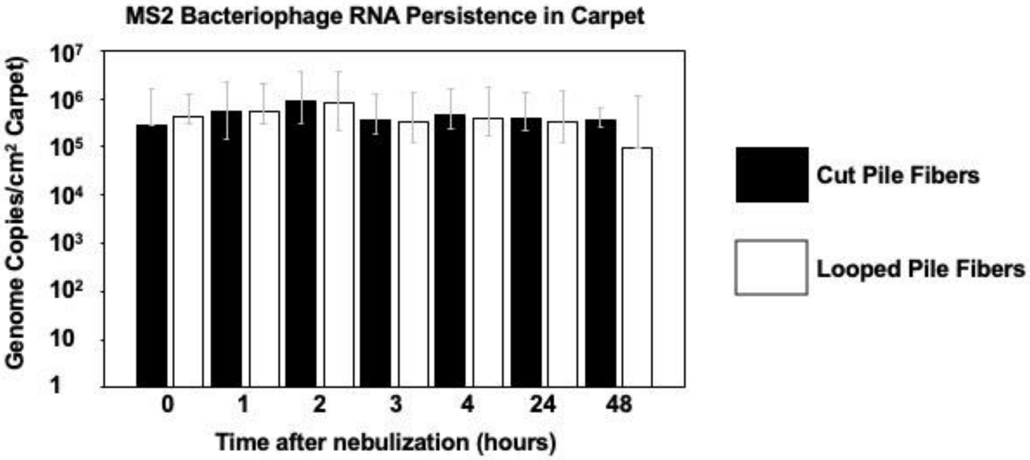

**Figure S7:** MS2 bacteriophage RNA concentrations in cut and looped carpet fibers. Error bars represent 95% confidence intervals.

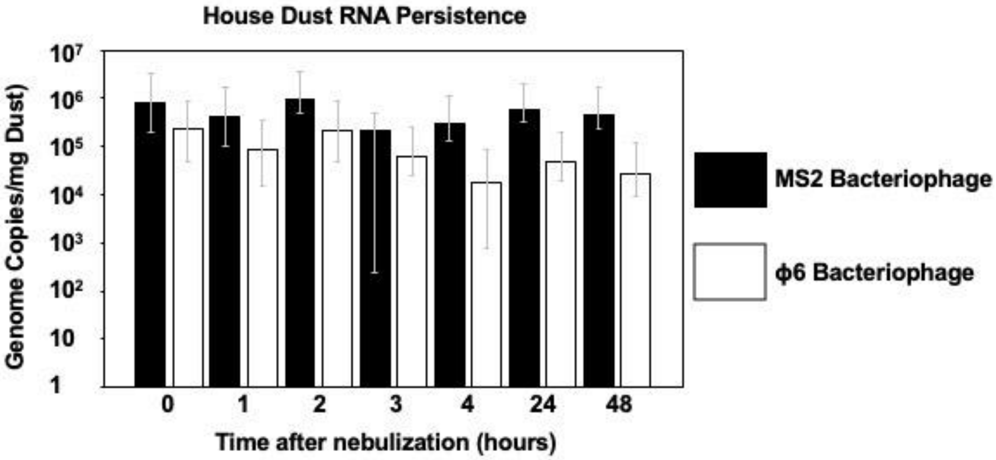

**Figure S8:** MS2 and Phi6 bacteriophage RNA concentrations in house dust. Error bars represent 95% confidence intervals.

**Table S1**:Modified artificial saliva recipe using 1000 mL of DI water (Woo et al. 2010).

| **Reagent** | **Quantity** |
|---|---|
| $MgCl_2 \cdot 6\ H_2O$ | 0.04 g |
| $CaCl_2 \cdot 2\ H_2O$ | 0.13 g |
| $NaHCO_3$ | 0.42 g |
| 0.2 M $KH_2PO_4$ | 7.70 mL |
| 0.2 M $K_2HPO_4$ | 12.3 mL |
| $NH_4Cl$ | 0.11 g |
| KSCN | 0.19 g |
| $(NH_2)_2CO$ | 0.12 g |
| NaCl | 0.88 g |
| KCl | 1.04 g |
| Gastric mucin | 3.00 g |
| Distilled water | 1000 mL |
| pH | 7 |

**Table S2:** First-order decay rate constants for Phi6 and MS2 bacteriophages in artificial saliva.

| Phage Viability | Conditions | k ($hr^{-1}$) | CI | $R^2$ | $T_{90}$ (hrs) | $T_{99}$ (hrs) |
|---|---|---|---|---|---|---|
| Phi6 | saliva | -0.04 | -0.07 to -0.01 | 0.60 | 60.61 | 121.21 |
| MS2 | saliva | 0.07 | 0.05 to 0.08 | 0.96 | N/A | |

N/A = Not applicable due to positive k

**Saliva Viability**

**MS2 Bacteriophage**

| Time Point (hrs) | Viability (PFU/mL) |
|---|---|
| 0 | 300000 |
| 1 | 167000 |
| 2 | 333000 |
| 3 | 467000 |
| 4 | 433000 |
| 48 | 7000000 |

**PHI6 Bacteriophage**

| Time Point (hrs) | Viability (PFU/mL) |
|---|---|
| 0 | 167000000 |
| 1 | 200000000 |
| 2 | 100000000 |
| 3 | 133000000 |
| 4 | 40000000 |
| 48 | 30000000 |

**Table S3:** Phi6 bacteriophage viability values for cut and looped carpet fibers (PFU/cm$^2$).

**Phi6 Bacteriophage Carpet Viability**

| Time Point (hrs) | Carpet Type | Viability (PFU/cm^2 carpet) |
|---|---|---|
| 0 | Cut | 34400 |
| | Cut | 12444 |
| | Cut | 16889 |
| 1 | Cut | 6489 |
| | Cut | 1907 |
| | Cut | 10560 |
| 2 | Cut | 313 |
| | Cut | 338 |
| | Cut | 422 |
| 3 | Cut | 44 |
| | Cut | 185 |
| | Cut | N.D. |
| 4 | Cut | 63 |
| | Cut | 36 |
| | Cut | 50 |
| 24 | Cut | N.D. |
| | Cut | N.D. |
| | Cut | N.D. |
| 48 | Cut | N.D. |
| | Cut | N.D. |
| | Cut | N.D. |

| Time Point (hrs) | Carpet Type | Viability (PFU/cm^2 carpet) |
|---|---|---|
| 0 | Loop | 3467 |
| | Loop | 12723 |
| | Loop | 9426 |
| 1 | Loop | 11400 |
| | Loop | 2848 |
| | Loop | 1044 |
| 2 | Loop | 6222 |
| | Loop | 622 |
| | Loop | 62 |
| 3 | Loop | 1111 |
| | Loop | 10 |
| | Loop | N.D. |
| 4 | Loop | 256 |
| | Loop | 256 |
| | Loop | 220 |
| 24 | Loop | 8 |
| | Loop | 29 |
| | Loop | 10 |
| 48 | Loop | N.D. |
| | Loop | N.D. |
| | Loop | N.D. |

***N.D. refers to non-detectable or below detection limit (6 PFU/cm$^2$ carpet)**

**Table S4:** MS2 bacteriophage viability values for cut and looped carpet fibers (PFU/cm$^2$).

**MS2 Bacteriophage Carpet Viability**

| Time Point (hrs) | Carpet Type | Viability (PFU/cm² carpet) | Time Point (hrs) | Carpet Type | Viability (PFU/cm² carpet) |
|---|---|---|---|---|---|
| 0 | Cut | 8409 | 0 | Loop | 1173 |
| | Cut | 11822 | | Loop | 474 |
| | Cut | 7431 | | Loop | 1540 |
| 1 | Cut | 1233 | 1 | Loop | 777 |
| | Cut | 1032 | | Loop | 896 |
| | Cut | 1272 | | Loop | 773 |
| 2 | Cut | 1434 | 2 | Loop | 266 |
| | Cut | 845 | | Loop | 1182 |
| | Cut | 1971 | | Loop | 620 |
| 3 | Cut | 26 | 3 | Loop | 307 |
| | Cut | 56 | | Loop | 34 |
| | Cut | 10 | | Loop | 120 |
| 4 | Cut | 45 | 4 | Loop | 256 |
| | Cut | 127 | | Loop | 43 |
| | Cut | 20 | | Loop | 77 |
| 24 | Cut | 33 | 24 | Loop | 164 |
| | Cut | 93 | | Loop | 149 |
| | Cut | 50 | | Loop | 159 |
| 48 | Cut | N.D. | 48 | Loop | N.D. |
| | Cut | 17 | | Loop | 10 |
| | Cut | N.D. | | Loop | N.D. |

***N.D. refers to non-detectable or below detection limit (6 PFU/cm² carpet)**

**Table S5:** MS2 and Phi6 bacteriophage viability values house dust (PFU/mg).

**Dust Viability**

**MS2 Bacteriophage**

| Time Point (hrs) | Viability (PFU/mg House Dust) |
|---|---|
| 0 | 1008 |
| | 6266 |
| | 1204 |
| 1 | 298 |
| | 2507 |
| | 379 |
| 2 | 293 |
| | 139 |
| | 114 |
| 3 | 12 |
| | 17 |
| | 29 |
| 4 | 249 |
| | 25 |
| | 206 |
| 24 | 520 |
| | 548 |
| | 300 |
| 48 | 12 |
| | 24 |
| | 13 |

**PHI6 Bacteriophage**

| Time Point (hrs) | Viability (PFU/mg House Dust) |
|---|---|
| 0 | 824 |
| | 7040 |
| | 984 |
| 1 | 598 |
| | N.D. |
| | 13 |
| 2 | 2 |
| | 1 |
| | 1 |
| 3 | N.D. |
| | N.D. |
| | N.D. |
| 4 | N.D. |
| | N.D. |
| | N.D. |
| 24 | N.D. |
| | N.D. |
| | N.D. |
| 48 | N.D. |
| | N.D. |
| | N.D. |

***N.D. refers to non-detectable or below detection limit (6 PFU/mg Dust)**

**Table S6:** Phi6 bacteriophage RNA values for cut and looped carpet (Genome Copies/cm$^2$ carpet).

**Phi6 Bacteriophage Carpet RNA qPCR Values**

| Time Point (hrs) | Carpet Type | RNA (Genome Copies/cm² carpet) | Time Point (hrs) | Carpet Type | RNA (Genome Copies/cm² carpet) |
|---|---|---|---|---|---|
| 0 | Cut | 173032 | 0 | Loop | 90141 |
| | Cut | 220105 | | Loop | 27808 |
| | Cut | 178014 | | Loop | 234113 |
| | Cut | 298739 | | Loop | 250854 |
| | Cut | 330899 | | Loop | 282934 |
| | Cut | 268343 | | Loop | 276572 |
| | Cut | 227867 | | Loop | N.D. |
| | Cut | 107474 | | Loop | N.D. |
| | Cut | 107757 | | Loop | 929061 |
| 1 | Cut | 395095 | 1 | Loop | N.D. |
| | Cut | 151001 | | Loop | N.D. |
| | Cut | 331667 | | Loop | 84081 |
| | Cut | 103315 | | Loop | 2707844 |
| | Cut | 111356 | | Loop | 1959389 |
| | Cut | 144426 | | Loop | 2105038 |
| | Cut | N.D. | | Loop | 306488 |
| | Cut | N.D. | | Loop | 176519 |
| | Cut | N.D. | | Loop | 351594 |
| 2 | Cut | 800719 | 2 | Loop | 632780 |
| | Cut | 584146 | | Loop | 547205 |
| | Cut | 578752 | | Loop | 671751 |
| | Cut | 468254 | | Loop | 1531442 |
| | Cut | 287549 | | Loop | 1763782 |
| | Cut | 136643 | | Loop | 1510268 |
| | Cut | 994435 | | Loop | 592513 |
| | Cut | 1115876 | | Loop | 793044 |
| | Cut | 1890688 | | Loop | 342970 |
| 3 | Cut | 1071979 | 3 | Loop | 64906 |
| | Cut | 891548 | | Loop | 158149 |
| | Cut | 1479184 | | Loop | 88545 |
| | Cut | 1988793 | | Loop | N.D. |

| | | | | | |
|---|---|---|---|---|---|
| | Cut | 3110798 | | Loop | N.D. |
| | Cut | 2782139 | | Loop | 310016 |
| | Cut | 909786 | | Loop | 355948 |
| | Cut | 707770 | | Loop | 201487 |
| | Cut | 671183 | | Loop | 100527 |
| 4 | Cut | 1013461 | 4 | Loop | 359054 |
| | Cut | 1498656 | | Loop | 180166 |
| | Cut | 1694914 | | Loop | 370567 |
| | Cut | 2439294 | | Loop | 166724 |
| | Cut | 1578688 | | Loop | 100285 |
| | Cut | 1885390 | | Loop | N.D. |
| | Cut | 1340522 | | Loop | 130043 |
| | Cut | 723530 | | Loop | 65167 |
| | Cut | 616705 | | Loop | 174730 |
| 24 | Cut | N.D. | 24 | Loop | 210573 |
| | Cut | N.D. | | Loop | 141877 |
| | Cut | N.D. | | Loop | 329473 |
| | Cut | N.D. | | Loop | 160608 |
| | Cut | 890904 | | Loop | 353959 |
| | Cut | 714503 | | Loop | 182411 |
| | Cut | 379216 | | Loop | 130835 |
| | Cut | 495753 | | Loop | 340795 |
| | Cut | 91293 | | Loop | 406108 |
| 48 | Cut | 111804 | 48 | Loop | 85748 |
| | Cut | 105994 | | Loop | 68701 |
| | Cut | 199818 | | Loop | 26234 |
| | Cut | N.D. | | Loop | 89134 |
| | Cut | 1060288 | | Loop | 159168 |
| | Cut | 1430496 | | Loop | N.D. |
| | Cut | 239428 | | Loop | 33035 |
| | Cut | 338310 | | Loop | N.D. |
| | Cut | 417232 | | Loop | 28302 |

***N.D. refers to non-detectable or below detection limit for qPCR**

**Table S7:** MS2 bacteriophage RNA values for cut and looped carpet (Genome Copies/$cm^2$ carpet).

**MS2 Bacteriophage Carpet RNA qPCR Values**

| Time Point (hrs) | Carpet Type | RNA (Genome Copies/$cm^2$ carpet) | Time Point (hrs) | Carpet Type | RNA (Genome Copies/$cm^2$ carpet) |
|---|---|---|---|---|---|
| 0 | Cut | 458954 | 0 | Loop | 105742 |
| | Cut | 390134 | | Loop | 73603 |
| | Cut | 298430 | | Loop | 3172 |
| | Cut | 240315 | | Loop | 286055 |
| | Cut | 246995 | | Loop | 292114 |
| | Cut | 234234 | | Loop | 352728 |
| | Cut | 173134 | | Loop | 766767 |
| | Cut | 174698 | | Loop | 819571 |
| | Cut | 198584 | | Loop | 1219540 |
| 1 | Cut | 526271 | 1 | Loop | 322842 |
| | Cut | 336260 | | Loop | 348124 |
| | Cut | 482727 | | Loop | 280175 |
| | Cut | 466474 | | Loop | 1218555 |
| | Cut | 522001 | | Loop | 1414011 |
| | Cut | 473778 | | Loop | 737820 |
| | Cut | 810159 | | Loop | 204561 |
| | Cut | 654698 | | Loop | 149022 |
| | Cut | 636648 | | Loop | 226464 |
| 2 | Cut | 785180 | 2 | Loop | 292280 |
| | Cut | 509468 | | Loop | 801394 |
| | Cut | 503890 | | Loop | 576686 |
| | Cut | 601621 | | Loop | 902078 |
| | Cut | 624104 | | Loop | 1515643 |
| | Cut | 412062 | | Loop | 1046591 |
| | Cut | 1493547 | | Loop | 55717 |
| | Cut | 1603943 | | Loop | 993047 |
| | Cut | 1668199 | | Loop | 947819 |
| 3 | Cut | 471229 | 3 | Loop | 324570 |
| | Cut | 305540 | | Loop | 552936 |
| | Cut | 337596 | | Loop | 451988 |
| | Cut | 235688 | | Loop | 303836 |

| | | |
|---|---|---|
| | Cut | 689080 |
| | Cut | 254032 |
| | Cut | 422083 |
| | Cut | 201506 |
| | Cut | 215197 |
| 4 | Cut | 377607 |
| | Cut | 429421 |
| | Cut | 550903 |
| | Cut | 620222 |
| | Cut | 674071 |
| | Cut | 396220 |
| | Cut | 323414 |
| | Cut | 461223 |
| | Cut | 361966 |
| 24 | Cut | 351173 |
| | Cut | 298348 |
| | Cut | 234053 |
| | Cut | 575996 |
| | Cut | 641199 |
| | Cut | 448916 |
| | Cut | 399443 |
| | Cut | 232590 |
| | Cut | 293899 |
| 48 | Cut | 148366 |
| | Cut | 166782 |
| | Cut | 133007 |
| | Cut | 627321 |
| | Cut | 448961 |
| | Cut | 644214 |
| | Cut | 278232 |
| | Cut | 323919 |
| | Cut | 346141 |

| | | |
|---|---|---|
| | Loop | 236049 |
| | Loop | 322234 |
| | Loop | 194835 |
| | Loop | 256100 |
| | Loop | 193007 |
| 4 | Loop | 657044 |
| | Loop | 676257 |
| | Loop | 189214 |
| | Loop | 370668 |
| | Loop | 252416 |
| | Loop | 361949 |
| | Loop | 136689 |
| | Loop | 356442 |
| | Loop | 408930 |
| 24 | Loop | 281999 |
| | Loop | 320897 |
| | Loop | 175340 |
| | Loop | 288731 |
| | Loop | 202440 |
| | Loop | 287660 |
| | Loop | 447029 |
| | Loop | 474548 |
| | Loop | 463057 |
| 48 | Loop | 197924 |
| | Loop | 150176 |
| | Loop | 216212 |
| | Loop | 5094 |
| | Loop | 3560 |
| | Loop | 8427 |
| | Loop | 92134 |
| | Loop | 64225 |
| | Loop | 80213 |

**Table S8:** MS2 and Phi6 bacteriophage RNA values for house dust (Genome Copies/mg dust

**House Dust RNA qPCR Values**

| MS2 Bacteriophage | | | Phi6 Bacteriophage | |
|---|---|---|---|---|
| **Time Point (hrs)** | **RNA (Genome Copies/mg Dust)** | | **Time Point (hrs)** | **RNA (Genome Copies/mg Dust)** |
| 0 | 358074 | | 0 | 125167 |
| | 230754 | | | 80461 |
| | 206418 | | | 86026 |
| | 1690840 | | | 738140 |
| | 1757282 | | | 438840 |
| | 1684321 | | | 349109 |
| | 417342 | | | 101899 |
| | 394932 | | | 40583 |
| | 390632 | | | 86996 |
| 1 | 246754 | | 1 | 73112 |
| | 256237 | | | 35945 |
| | 271714 | | | 59709 |
| | 880647 | | | 217994 |
| | 1061961 | | | 180949 |
| | 729835 | | | 194070 |
| | 115850 | | | 18018 |
| | 113567 | | | 11318 |
| | 122474 | | | 12017 |
| 2 | 687750 | | 2 | 118463 |
| | 698570 | | | 68819 |
| | 719931 | | | 103180 |
| | 809799 | | | 78463 |
| | 926351 | | | 58821 |
| | 718362 | | | 78267 |
| | 1074150 | | | 598255 |
| | 1677437 | | | 322029 |
| | 1075584 | | | 499392 |
| 3 | 204759 | | 3 | 41422 |
| | 107712 | | | 32039 |
| | 114225 | | | 38550 |
| | N.D. | | | 55671 |
| | N.D. | | | 36034 |

| | | | |
|---|---|---|---|
| | N.D. | | 62066 |
| | 686710 | | 107333 |
| | 341304 | | 79813 |
| | 434458 | | 98896 |
| 4 | 186281 | 4 | 21915 |
| | 286695 | | 14408 |
| | 225770 | | 20964 |
| | 195519 | | 28611 |
| | 204031 | | 22591 |
| | 206042 | | 9370 |
| | 426525 | | 40237 |
| | 436824 | | 7036 |
| | 450947 | | N.D. |
| 24 | 714314 | 24 | 56433 |
| | 762156 | | 36987 |
| | 588096 | | 103150 |
| | 585788 | | 69554 |
| | 636722 | | 53865 |
| | 404650 | | 18281 |
| | 539639 | | 46805 |
| | 435681 | | 27361 |
| | 429315 | | 33545 |
| 48 | 491668 | 48 | 51338 |
| | 532173 | | 19076 |
| | 345959 | | 25944 |
| | 404220 | | 38362 |
| | 266795 | | 4657 |
| | 380971 | | 23094 |
| | 592296 | | 33378 |
| | 416687 | | 18307 |
| | 543369 | | 39317 |

***N.D. refers to non-detectable or below detection limit for qPCR**

**Table S9:** Phi6 and MS2 bacteriophage viability values for untreated (no cleaning), vacuumed, hot water extraction, and steam cleaning methods for cut carpet fibers. (PFU/cm$^2$).

**Cleaning Methods: Viability**

**MS2 Bacteriophage**

| EXPERIMENT 1 | | |
|---|---|---|
| CLEANING METHOD | Carpet type | Viability (PFU/cm^2 carpet) |
| STEAM CLEAN | Cut | N.D. |
| | Cut | N.D. |
| | Cut | N.D. |
| VACUUM | Cut | 240 |
| | Cut | 235 |
| | Cut | 106 |
| NO CLEAN | Cut | 249 |
| | Cut | 831 |

**PHI6 Bacteriophage**

| EXPERIMENT 1 | | |
|---|---|---|
| CLEANING METHOD | Carpet type | Viability (PFU/cm^2 carpet) |
| STEAM CLEAN | Cut | N.D. |
| | Cut | N.D. |
| | Cut | N.D. |
| VACUUM | Cut | N.D. |
| | Cut | N.D. |
| | Cut | 21 |
| NO CLEAN | Cut | 4907 |
| | Cut | 521 |

**MS2 Bacteriophage**

| EXPERIMENT 2 | | |
|---|---|---|
| CLEANING METHOD | Carpet type | Viability (PFU/cm^2 carpet) |
| Disinfectant (PURTABS) | Cut | N.D. |
| | Cut | N.D. |
| | Cut | N.D. |
| Hot Water Extraction | Cut | 117 |
| | Cut | N.D. |
| | Cut | 365 |
| NO CLEAN | Cut | 411 |

**PHI6 Bacteriophage**

| EXPERIMENT 2 | | |
|---|---|---|
| CLEANING METHOD | Carpet type | Viability (PFU/cm^2 carpet) |
| Disinfectant (PURTABS) | Cut | N.D. |
| | Cut | N.D. |
| | Cut | N.D. |
| Hot Water Extraction | Cut | 17 |
| | Cut | N.D. |
| | Cut | 7 |
| NO CLEAN | Cut | 24 |

***N.D. refers to non-detectable or below detection limit (6 PFU/cm$^2$ carpet)**

**Table S10:** Phi6 and MS2 bacteriophage RNA values for untreated (no cleaning), vacuumed, hot water extraction, and steam cleaning methods for cut carpet fibers. (PFU/$cm^2$).

**Cleaning Methods RNA Persistence**

| MS2 Bacteriophage | | | PHI6 Bacteriophage | | |
|---|---|---|---|---|---|
| **CLEANING METHOD** | **Carpet type** | **RNA (Genome Copies/cm^2 carpet)** | **CLEANING METHOD** | **Carpet type** | **RNA (Genome Copies/cm^2 carpet)** |
| STEAM CLEAN | Cut | 69912 | STEAM CLEAN | Cut | 1111929 |
| | Cut | 87515 | | Cut | 607330 |
| | Cut | 107806 | | Cut | 484558 |
| | Cut | 157630 | | Cut | 391419 |
| | Cut | 131908 | | Cut | 1033254 |
| | Cut | 91079 | | Cut | 888537 |
| | Cut | 88615 | | Cut | 323537 |
| | Cut | 95627 | | Cut | 269678 |
| | Cut | 77437 | | Cut | 250831 |
| Vacuum | Cut | 57633 | Vacuum | Cut | 113846 |
| | Cut | 63825 | | Cut | 148957 |
| | Cut | 66597 | | Cut | 101379 |
| | Cut | 10846 | | Cut | 746708 |
| | Cut | 200091 | | Cut | 1266056 |
| | Cut | 214404 | | Cut | N.D |
| | Cut | 151761 | | Cut | 745693 |
| | Cut | 232444 | | Cut | 678956 |
| | Cut | 206788 | | | 1385252 |
| Disinfectant (PURTABS) | Cut | N.D | Disinfectant (PURTABS) | Cut | 44132 |
| | Cut | N.D | | Cut | 190426 |
| | Cut | N.D | | Cut | 68409 |
| | Cut | N.D | | Cut | 199562 |
| | Cut | N.D | | Cut | N.D |
| | Cut | N.D | | Cut | 62420 |
| | Cut | N.D | | Cut | 134660 |
| | Cut | N.D | | Cut | 82898 |
| | Cut | N.D | | Cut | 148235 |
| | Cut | 99224 | | Cut | N.D |

| | | | | | |
|---|---|---|---|---|---|
| **Hot Water Extraction** | **Cut** | **118894** | **Hot Water Extraction** | **Cut** | **141709** |
| | **Cut** | **90219** | | **Cut** | **113928** |
| | **Cut** | **49712** | | **Cut** | **68311** |
| | **Cut** | **46091** | | **Cut** | **N.D** |
| | **Cut** | **41492** | | **Cut** | **N.D** |
| | **Cut** | **62818** | | **Cut** | **111754** |
| | **Cut** | **79879** | | **Cut** | **113254** |
| | **Cut** | **85166** | | **Cut** | **179605** |
| **No Clean** | **Cut** | **99108** | **No Clean** | **Cut** | **N.D** |
| | **Cut** | **100315** | | **Cut** | **N.D** |
| | **Cut** | **98607** | | **Cut** | **446340** |
| | **Cut** | **138707** | | **Cut** | **922131** |
| | **Cut** | **111437** | | **Cut** | **740889** |
| | **Cut** | **148307** | | **Cut** | **895000** |
| | **Cut** | **489233** | | **Cut** | **476285** |
| | **Cut** | **491886** | | **Cut** | **947293** |
| | **Cut** | **479668** | | **Cut** | **389572** |
| **Negative Control** | **Cut** | **N.D** | **Negative Control** | **Cut** | **N.D** |
| | **Cut** | **N.D** | | **Cut** | **N.D** |
| | **Cut** | **N.D** | | **Cut** | **N.D** |

***N.D. refers to non-detectable or below detection limit for qPCR**
